# Retargeting: an unrecognized consideration in endonuclease-based gene drive biology

**DOI:** 10.1101/089946

**Authors:** Andrew M. Scharenberg, Barry L Stoddard, Raymond J Monnat, Anthony Nolan

## Abstract

There is intense interest surrounding the use of gene editing nucleases in gene drive systems to control agricultural insect pests and insect vectors of infectious diseases such as malaria, dengue and Zika virus. While gene drive systems offer immense promise for the beneficial modification of deleterious insect populations, their unique mechanism of action also raises novel safety concerns and regulatory issues. A recent US National Academies of Science report provides a list of potential regulatory issues associated with implementation of homologous recombination (HR)-mediated gene drive systems, based on the premise that all such systems would exhibit similar biological behaviors. Here we examine how HR-mediated gene drive systems based on different gene editing nuclease platforms could be affected by mutations that occur during host cell transcription, genome replication, and, in conjunction with gene editing nuclease activity, during HR-mediated gene drive. Our analysis suggests that the same feature that makes RNA-guided nucleases such attractive research tools—their ease of reprogramming by alterations to their guide RNA components—might also contribute to increased rates of retargeting that could influence the long term behavior of RNA-guided gene drive systems. Predictability of behavior over time is an issue that should be addressed by in-depth risk assessment before field testing of organisms incorporating nuclease-mediated gene drives.

---

Homologous-recombination (HR)-based gene drive systems aimed at population-wide modification of agricultural pest and insect disease vector populations (hereafter “HR-gene drives”) are among the most broadly impactful potential applications of gene editing technologies. HR-gene drives involve the propagation of heritable alleles encoding gene editing nucleases in a population of wild organisms, with the intent of modifying the reproductive potential or vector competence of that population. Given the self-propagating nature of gene drives, a systematic evaluation of potential safety issues is essential prior to any anticipated field release.

A recent US National Academies of Science (NAS) report has provided an analysis of safety and regulatory issues associated with the development of HR-mediated gene drives ^1^. While this NAS report represents a critical step forward in the development and eventual use of gene drive systems, the analyses presented were built on the assumption that different nuclease platforms used to enable gene drive would share similar biological behavior profiles. Here, we assess how known sources of mutation could alter the behavior of HR-gene drive systems that rely upon different types of gene editing nucleases. Our analysis highlights the potential susceptibility of RNA-guided gene drive systems (relative to protein nuclease-based gene drives) to mutational “re-targeting”.

All HR-based gene drive systems are designed to display the super-Mendelian inheritance characteristic of the life cycle of naturally occurring, invasive mobile introns (Figure 1). Invasive mobile introns are parasitic DNA elements that, following mating of their host organism, propagate by copying themselves from their genomic location to a homologous host allele. This form of gene transfer, referred to as ‘homing’, is targeted and initiated by the ability of an endonuclease encoded in the mobile intron to cleave genomic target sites that lack a copy of the mobile intron. The resulting process of HR-mediated double strand break repair of the cleaved genomic target site results in the copying of the mobile intron and its encoded endonuclease to create a new, intron-containing allele with the potential to further propagate. The ability of this transfer reaction to occur whenever a cleavage-sensitive intron-lacking allele is present accounts for the ability of mobile introns to achieve super-Mendelian frequencies of inheritance, with the potential to occupy essentially all empty host alleles in a naïve population after relatively few generations ^2^. This type of ‘smart DNA parasitism’ is further enabled by the ability of many mobile introns and their encoded endonucleases to target host genes, including essential genes, in a way that minimally perturbs the host gene function after intron insertion. This ‘invisibility cloak’ is typically conferred by the ability of mobile introns to self-splice at the RNA or protein level, thus avoiding elimination or imposing a fitness cost or genetic load on the host organism.

**Figure 1:**
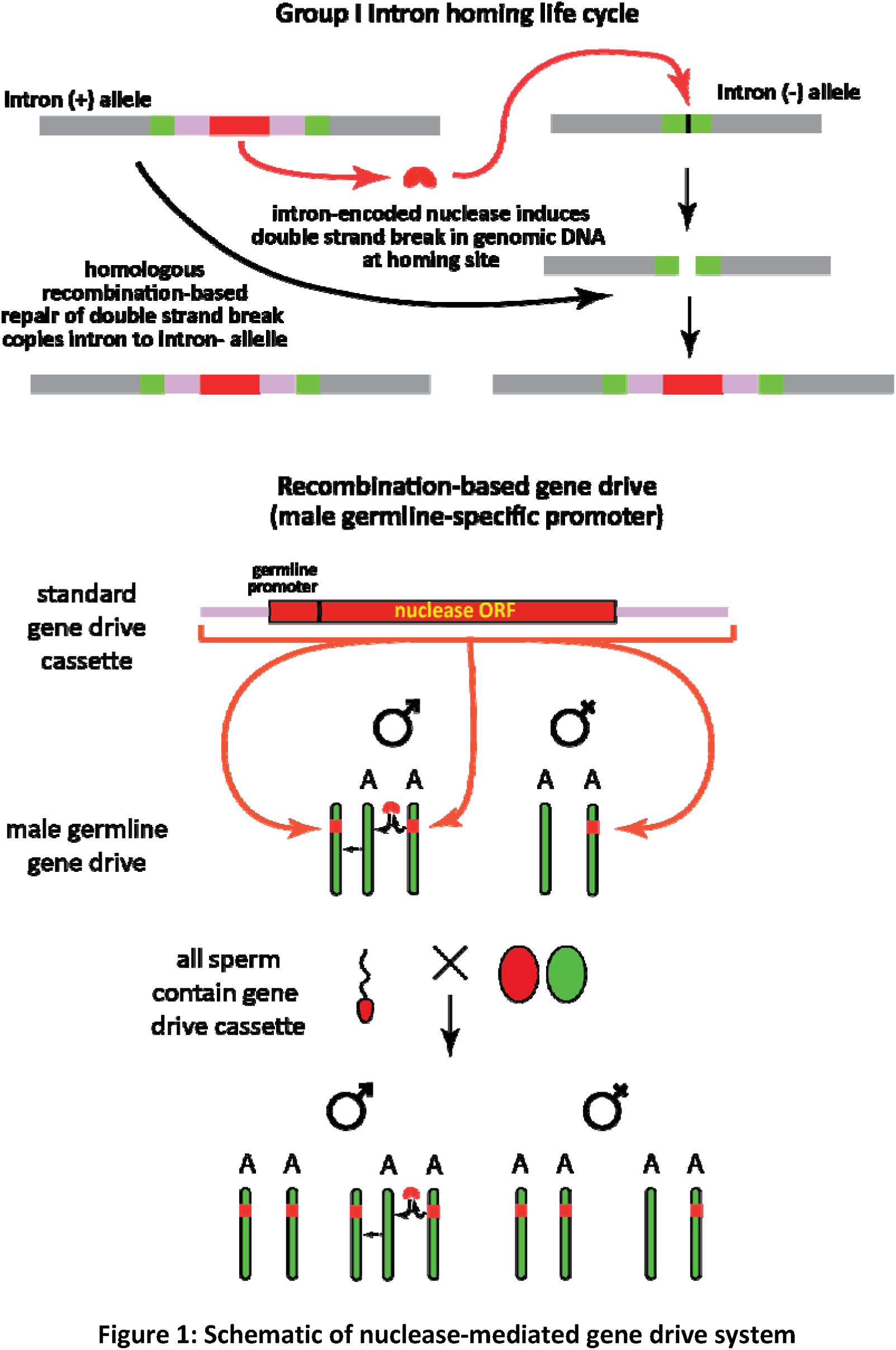
Schematic of nuclease-mediated gene drive system

The potential of synthetic HR-based gene drive systems to propagate in multi-cellular organisms was first described by Burt ^3^. The synthetic HR-gene drive systems he described are based on the same homologous recombination-based propagation mechanism as a natural mobile intron. However, synthetic HR-gene drive cassettes differ from natural mobile introns by encoding a nuclease engineered to cleave a conserved a target site in a host gene only when expressed in the germline, resulting in the HR-gene drive cassette copying itself (i.e. “homing”) into unmodified host gene alleles only in the germline. Burt and colleagues also described the potential application of HR-based gene drive systems to modify the fecundity of the target organism ^3^, or the competence of target organisms to serve as vectors of an infectious disease ^3^.

As HR-gene drives necessarily involve the modification of entire populations of wild organisms to encode and express self-propagating synthetic genes, potential biosafety issues were raised and considered very early on in the development of this technology. The recent NAS report represents an important formalization of the associated issues, and articulates a regulatory roadmap to guide further development of gene drive technologies for any application ^1^. The major biosafety issues identified with population-control gene drives were divided into two general categories which can be most readily understood and discussed by considering the consequences of drive as a function of genomic locus and host organism. Unintended alterations in function of a <u>target organism,</u> due to an unanticipated modification of its physiology, were termed “<u>off target</u>” impacts of a gene drive system. “Off target” impacts can be separated into unintended consequences of modifying the intended genomic locus (“on locus/off target” impact) or any other genomic locus (“off locus/off target” impact) in the intended host (target) organism. The possibility of hybridization followed by introgression of a gene drive cassette to an organism other than the intended host raises the additional consideration of potential “non-target impacts” such as propagation and population-level modification of the fecundity, or other aspects, of a non-target organism.

As summarized in the NAS report, a large body of data and experience is available to assess and potentially manage “off target” effects. Investigators involved in the development of gene drive technology have also proposed many approaches to mitigate such affects^3-6^. The NAS report also provides a formalization of potential “off-target” and “non-target” impacts, and correctly points out that the likelihood of these events is harder to predict, and thus it is potentially more difficult to develop corresponding, effective mitigation approaches for them.

While the NAS report is extremely comprehensive, we note an aspect of gene drive biology that was not considered: the potential of mutation to modify the DNA target sequence cleavage properties of a gene drive nuclease. As all DNA sequences in a genome are subject to mutational variation, mutations will inevitably occur in any gene drive cassette over time. In the case of an HR-gene drive, such mutations require particular consideration because they may modify DNA base pairs that determine the target recognition specificity of a gene drive nuclease leading to cleavage of an unintended target site, a phenomenon we refer to as “retargeting”. Retargeting would potentially lessen the predictability of behavior of an HR-gene drive system, as the susceptibility of its encoded nuclease to retargeting would be predicted to correlate with the capacity of the HR gene drive to modify or copy itself into alternative genomic loci.

The potential of mutations to alter target site recognition of a gene editing nuclease is reasonably inferred as being directly related to the ease with which a nuclease target site can be modified. Experience gained from the engineering of homing endonucleases has shown that individual random mutations typically lead to loss of function (reduced overall activity), and rarely directly to acquisition of a novel target specificity. Consequently, over the time-frame of gene drive through a population, homing endonucleases (or other “difficult to reprogram” nucleases such as zinc finger nucleases) would be predicted to be comparatively resistant to mutational retargeting: multiple mutations would be necessary to generate a variant enzyme with both efficient cleavage activity and a significantly altered cleavage specificity. TAL effector nucleases would be predicted to be somewhat more susceptible to retargeting, as individual mutations that alter the DNA binding residues of individual modular TAL repeats (termed ‘RVDs’) could lead to modified target specificity with relatively preserved cleavage activity. Perhaps of greater concern is the potential for mutational retargeting of RNA-guided nucleases (i.e. CRISPR/Cas9). Single mutations within the targeting sequence of a guide RNA in an RNA-guided gene drive would directly alter the cleavage specificity of the nuclease while leaving the nuclease fully active (Figure 2a). Similarly, insertion-deletion or translocation events that altered the targeting guide RNA sequence might lead to idiosyncratic retargeting of an RNA-guided gene drive system. These or other types of mutational alterations could also modify the lower stem loop of a guide RNA to allow the expressed nuclease to create double strand breaks in the guide (Figure 2a, e.g. as demonstrated in ^7^), increasing the probability of idiosyncratic retargeting. Thus the very feature that makes RNA-guided gene editing nucleases such popular research tools—the ease of target-specific engineering— would be predicted to increase the possibility of “off locus/off target” and “non-target” effects during RNA-guided endonuclease-driven gene drive. Some proposed safety mechanisms for RNA-guided nucleases such as “reversal” and “immunization” gene drive alleles would also be susceptible to this type of mutational or idiosyncratic retargeting ^4^. Recently described engineered Cas9 variants that display reduced cleavage activity at sequences closely related to that of their gRNA would be subject both to mutational alterations that alter the gRNA sequence, as well as mutations that revert backbone contact residues to wild-type, resulting in the reacquisition of the “off locus/off target” cleavage activity of the native enzyme.

A second important aspect of retargeting is the rate at which mutation would be predicted to affect different HR-gene drive architectures. All HR-gene drives are likely to be subjected to higher than background rates of mutation, as the DNA break-induced homologous recombination that underlies HR-gene drive propagation is thought to be error-prone relative to the mechanisms used in genome replication (the error rate DNA break-induced HR has been estimated at around 100-1000 times the background mutation rate in yeast^8^; similar data for animals will be important to develop). However, the choice of promoter may also influence the rate of mutation of a particular gene drive cassette, as higher transcriptional activity may be associated with higher rates of mutation due to transcription-associated mutation and recombination (^9-13^). This may be a particularly important consideration for RNA-guided gene drive cassette architectures that rely on high activity pol III promoters to drive guide RNA expression (Figure 2B).

**Figure 2 -.**
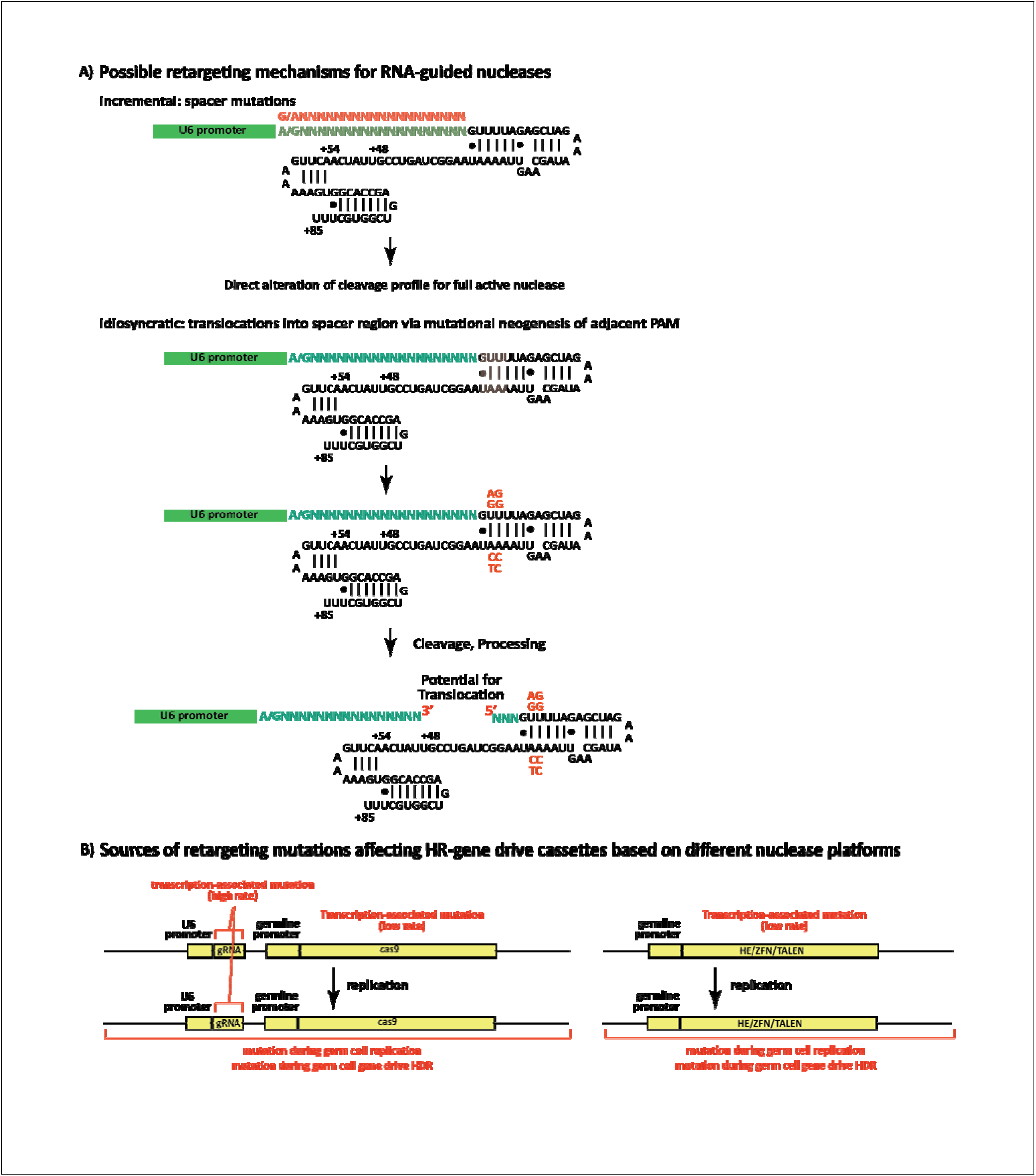
Mutational retargeting of gene drive systems

One example of how mutation-induced retargeting could be consequential is if the target species for a given gene drive has a closely related non-target species with which it regularly hybridizes to produce fertile offspring. Genomic and population level sequence data together with judicious choice of target site and careful nuclease engineering can be used to minimize the likelihood that a protein-encoded DNA recognition nuclease could initiate gene drive and spread through the non-target species after a hybridization event. This margin of safety may be diminished in the case of an RNA-guided drive system, where even careful target site selection could be undone by random point mutations within a guide RNA.

Given the potential for gene drive technologies to address serious public health issues, there is appropriate enthusiasm to continue their development, testing and eventual deployment. An important part of this effort will be to identify and minimize potential unintended consequences of any gene drive system by a thorough understanding of the underlying technology in the context of specific applications. The important differences in the biology of gene drive platforms, including the possibility described here that RNA-guided gene drive systems may be subject to mutational re-targeting at a higher rate than protein nuclease-guided gene drives, require thorough consideration in the risk assessment process recommended for any gene drive technology as these systems are developed and field trials are contemplated to address human parasitic and viral disease prevention ^1^.

## Acknowledgements

AMS, BLS, RJM, and TJN receive funding from the Target Malaria Consortium for their work on gene drive technology, including applications of homing endonucleases (AMS, BLS, RJM) and CRISPR/Cas9 (TJN) proteins as gene drive nucleases. AMS consults for and serves on the bluebird bio Scientific Advisory Board. He receives research support and, consulting income from bluebird bio, and holds equity in bluebird bio. BLS receives research support from bluebird bio, and holds equity in bluebird bio. RJM holds equity in bluebird bio.

